# GlpR Regulates Motility and Viscoelasticity Properties of *Pseudomonas aeruginosa*

**DOI:** 10.1101/2025.08.07.669157

**Authors:** Nicholas Evans, Hernan Grenett, Priscilla Grenett, Jessica A. Scoffield

## Abstract

Despite the onset of highly effective modulator therapies people with Cystic Fibrosis continue to experience recurrent microbial lung infections. Many of these individuals will have at least one positive culture per year for *Pseudomonas aeruginosa*, a bacterium that readily adapts to chronic CF airway disease. Of these adaptations, the formation of a protective biofilm and changes in motility are hallmarks of established infections. We have shown previous evidence that glycerol metabolism and the *P. aeruginosa* glycerol regulon repressor, GlpR, is linked to enhanced biofilm production and reduced susceptibility to tobramycin. In the current study, we report that loss of GlpR contributes to higher viscosity and elasticity in synthetic cystic fibrosis sputum media. Further, we show that the loss of the glycerol repressor GlpR, or growth on glycerol, both resulting in derepression of the GlpR/glycerol regulon, cause decreased motility in both acute and chronic CF-adapted lab strains. RNA sequencing analysis indicated that loss of GlpR altered the expression of genes involved in motility, iron scavenging, transport, metabolism, and virulence. An *in silico* search of *P. aeruginosa’s* genome using GlpR’s previously determined binding consensus site identified potential bindings sites in genes related to biofilm development, motility, antibiotic resistance, and metabolism, and these binding sites were confirmed using chromatin immunoprecipitation sequencing. Collectively, our results indicate that GlpR regulates *P. aeruginosa* phenotypes that facilitate persistence in the CF airway and we provide evidence that GlpR regulates genes outside of the canonical *glp* regulon.

**IMPORTANCE:** *Pseudomonas aeruginosa* continues to persist in the airways of individuals with cystic fibrosis (CF), even with modulator therapy. The nutritional environment of the CF airway has been shown to trigger the microevolution of *P. aeruginosa* to assist this bacterium in adaptation and persistence. *P. aeruginosa* can liberate glycerol from lung surfactant to use as a nutritional source. Previous studies have shown that glycerol metabolic genes are constitutively expressed in *P. aeruginosa* isolates recovered from CF sputum, highlighting the importance of the *glp* (glycerol) regulon, which is regulated by the transcriptional repressor, GlpR. Since glycerol is a critical nutritional source for *P. aeruginosa* adaptation, it is essential to understand the regulatory network controlled by GlpR.

## INTRODUCTION

Cystic fibrosis is a multi-system disease caused by mutations in the cystic fibrosis transmembrane conductance regulator channel (CFTR) protein (1). As a chloride channel, CFTR regulates mucus clearance and airway hydration, which are critical for the maintenance of a healthy airway. Mutations in CFTR inhibit proper mucociliary clearance, promote a dehydrated airway, and facilitate an accumulation of mucus that supports the colonization of microbes (1). Airway infections are typically multispecies and can persist for decades, ultimately leading to a loss of lung function (2–10).

*Pseudomonas aeruginosa* is major contributor of lung dysfunction and mortality in people with cystic fibrosis (pwCF). During infection in the CF airway, *P. aeruginosa* undergoes microevolution to adapt and persist in the lung (11–15). Genetic alterations that support the persistence of *P. aeruginosa* during chronic CF infections include the overproduction of alginate, loss of motility, and the redirection central metabolic processes to reprogram the expression of virulence genes or acquire nutrients. Airway sputum supports the growth and nutritional requirements of *P. aeruginosa* during CF infection (16–18). CF sputum is composed of both host-derived and bacterial-related products, including fatty acids, amino acids, acetate, and the major lung surfactant, phosphatidylcholine (PC) (16–19). Phospholipase C produced by *P. aeruginosa* cleaves phosphatidylcholine into phosphorylcholine, fatty acids, and glycerol, and as a result, liberated glycerol can be used as a potential nutritional source for *P. aeruginosa* (20–22). Previous studies have demonstrated that genes (*glpD* and *glpK*) specific to the *glp* (glycerol) regulon are constitutively expressed in some *P. aeruginosa* CF isolates recovered from sputum and these genes are controlled by the *glp* regulon repressor, GlpR (23, 24). Constitutive expression of genes involved in glycerol metabolism is likely due to the availability of host-derived glycerol in the lung which can serve as a nutrient source to fuel *P. aeruginosa* growth.

Previous studies by our group examining the role of the GlpR repressor in *P. aeruginosa* pathogenesis revealed that loss of GlpR promotes antibiotic tolerance, biofilm development, and alters the expression of virulence determinants (25, 26). Due to the relationship between biofilm development and a loss of motility, we questioned whether GlpR modulates adherence mechanisms that influence the biofilm mode of growth and clearance, including motility and viscoelasticity properties. Additionally, our previous work indicates that GlpR may regulate genes outside of the canonical *glp* regulon. Due to the importance of glycerol metabolism for *P. aeruginosa* adaptation in the CF airway and the likelihood that GlpR regulates non-*glp* genes, we sought to explore the role of GlpR on the *P. aeruginosa* transcriptome in CF and non-CF adapted isolates and identify the complete GlpR regulon. In this study, we report that loss of GlpR reduces motility, enhances viscoelasticity, and controls the *P. aeruginosa* transcriptome that influences biofilm development, motility, iron scavenging, and additional metabolic processes that are contribute to persistence. Further, using chromatin immunoprecipitation, we identified over 100 potential GlpR binding sites, thus providing evidence that GlpR regulates genes beyond the canonical *glp* regulon. In summary, our study reveals that GlpR regulates persistence phenotypes in the CF airway and controls an expanded regulon beyond the glycerol metabolic genes.

## RESULTS

### Loss of GlpR alters the viscoelasticity properties of CF-adapted *P. aeruginosa*

Glycerol is a readily available nutritional source for *P. aeruginosa* during colonization of the CF airway (22). We have previously demonstrated that the *glp* (glycerol) regulon repressor, GlpR, mediates biofilm development, antibiotic tolerance, and the overproduction of the biofilm exopolysaccharide alginate in the CF-adapted *P. aeruginosa* isolate FRD1 (25, 26). While investigating these phenotypes, we also observed that cultures of FRD1 and the FRD1 *ΔglpR* mutant in synthetic CF sputum (SCFM2) displayed differences in their viscosities, whereas the FRD1 Δ*glpR* mutant cultures were thicker and displayed a slower movement of fluid compared to wildtype FRD1 (Figures 1A and 1B). To quantify these observations, we utilized a rheometer to measure the viscoelastic properties of wildtype FRD1 and FRD1 Δ*glpR* mutant cultures grown in SCFM2. We found that across the range of applied shear rates, the FRD1 *ΔglpR* mutant was more viscous than FRD1 (Figure 1C). At lower shear rates, shear thinning occurred earlier for FRD1 than the FRD1 *ΔglpR* mutant as their peak viscosities were approximately one log apart (Figure 1D). We also detected a higher elasticity of the FRD1 *ΔglpR* mutant cultures compared to FRD1, indicating robust changes in their physical dynamics (Figure 1E).

**Figure 1.**
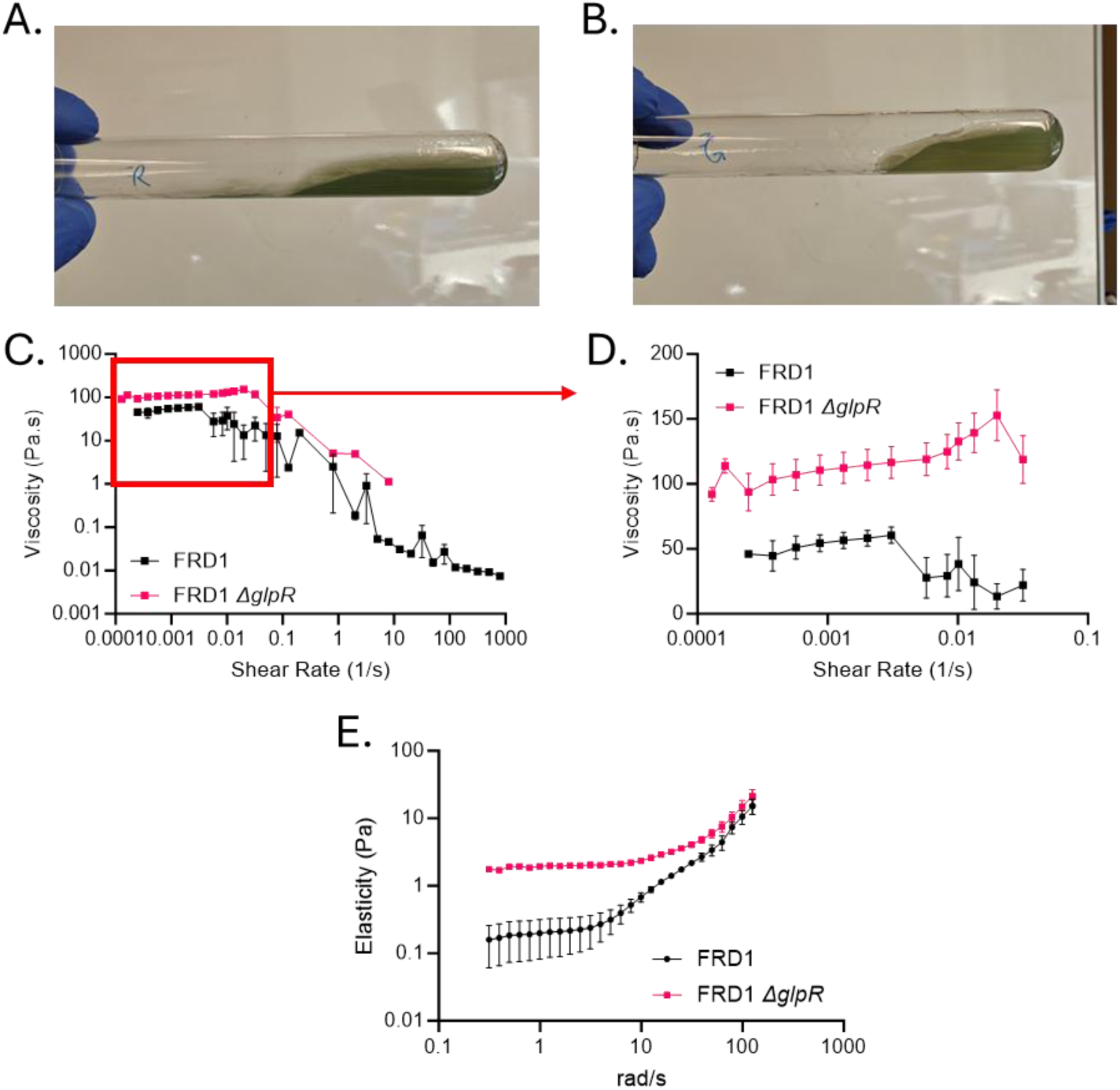
Loss of GlpR increases the viscosity and elasticity of cultures grown in SCFM2 culture. A) Image of spent culture of SCFM2 with FRD1 B) Image of spent SCFM2 culture with FRD1 *ΔglpR.* C) Rheometer measurements of the viscosity of spent SCFM2 with FRD1 and FRD1 glpR with increasing shear rates D) A graph of the lower shear rate range of the viscosity measurements FRD1 and FRD1 *ΔglpR* as measured by rheometer. E) A graph for the measured elasticity of the FRD1 and FRD1 *ΔglpR* grown in SCFM2. (n=3)

Based on our previous results showing increased alginate production by the FRD1 *ΔglpR* mutant (27), we hypothesized that alginate could be contributing to the changes we observed in the rheology of the bacterial cultures. To determine if alginate played a role, we added 10 U/mL of alginate lyase (AlgL), an enzyme that cleaves the glycosidic bonds of alginate, to the starting cultures. With AlgL supplementation, the rheometer no longer detected enough torque at lower shear rates for viscosity of FRD1 and FRD1 *ΔglpR*, with the AlgL-treated *glpR* mutant displaying similar viscosity to wildtype FRD1 (Figure 2A). Similarly, the addition of AlgL abolished the elasticity of FRD1 and FRD1 *ΔglpR* mutant (Figure 2B) cultures grown in SCFM2. These data indicate that loss of GlpR promotes alginate-dependent increased elasticity and viscosity in the CF-adapted isolate, FRD1.

**Figure 2.**
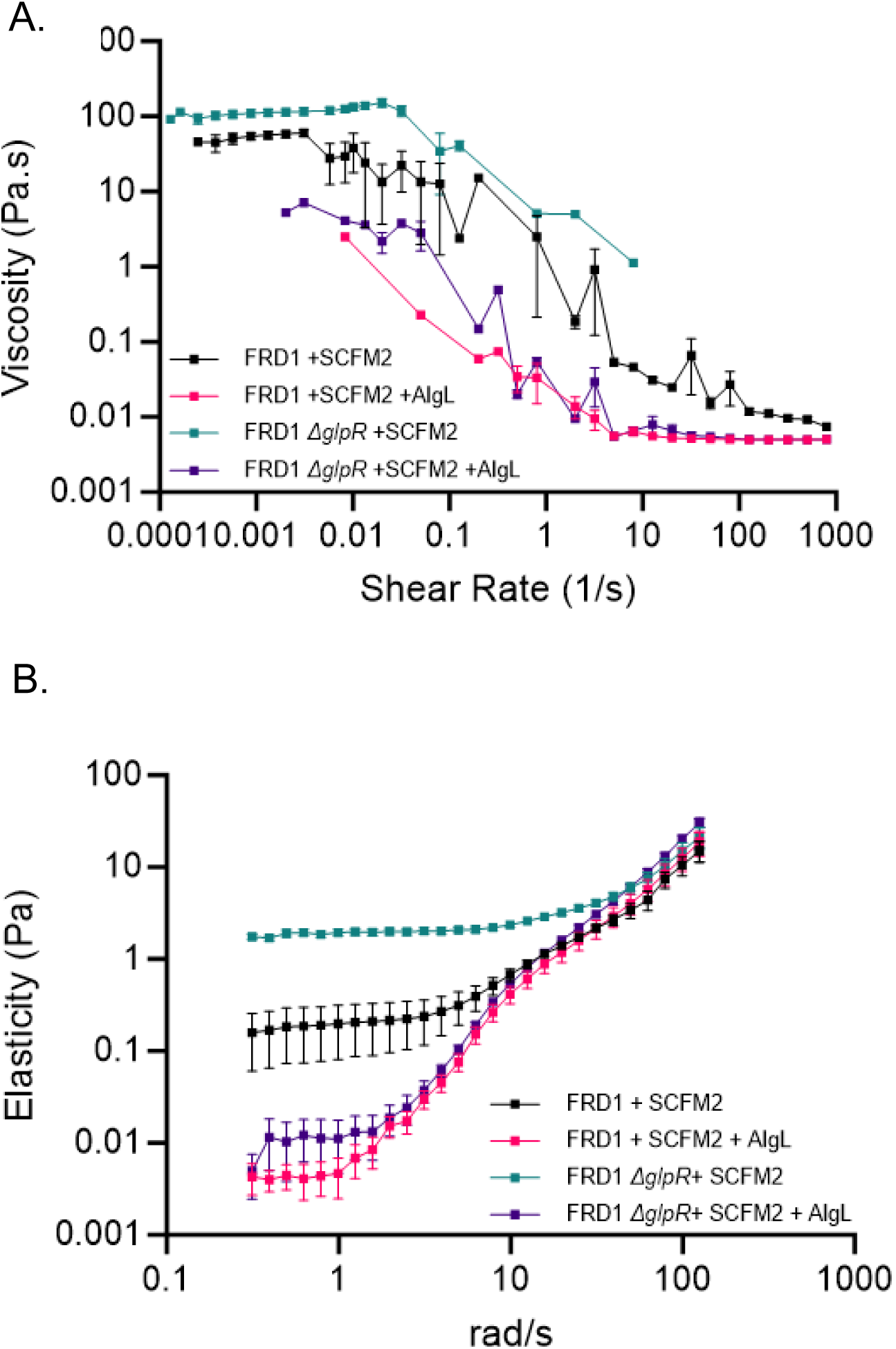
Alginate lyase abolishes increased viscosity and elasticity of FRD1 Δ*glpR* through the addition of alginate lyase. A) Viscosity measurements FRD1 and FRD1 *ΔglpR* grown for 16 hours in SCFM2 with and without alginate lyase treatment. B) Elasticity measurements FRD1 and FRD1 *ΔglpR* grown for 16 hours in SCFM2 with and without alginate lyase treatment. (n=3)

### Motility is reduced in GlpR-deficient mutants of *P. aeruginosa*

Our lab previously observed an increase in biofilm production on both glycerol and with the loss of GlpR (25). Further, it is well known that *P. aeruginosa* as well as other bacteria reduce motility during biofilm attachment (28–30). With this knowledge, we hypothesized that loss of GlpR reduces *P. aeruginosa* motility, hence we measured swarming, swimming, and twitching motility in the CF-adapted FRD1 strain and non-CF adapted PAO1 strain. As predicted, the PAO1 Δ*glpR* mutants exhibited reduced swimming, swarming, and twitching motility compared to wildtype PAO1 (Figures 3A - 3D). While the appearances of most strains were similar without GlpR despite the decreased size, we observed differences in morphology between PAO1 and the PAO1 *ΔglpR* mutant on the swarming plates. Most of the PAO1 swarms produced a more jagged or petal shaped margin with striations in the larger colony, whereas PAO1 *ΔglpR* had a smoother margin (Figure 3A). Loss of *glpR* reduced swimming and twitching motility in the FRD1 background compared to wildtype FRD1, but swarming motility was not significantly reduced (Figures 4A, 4E–4G). Complementation of *glpR* restored all defects in motility.

**Figure 3.**
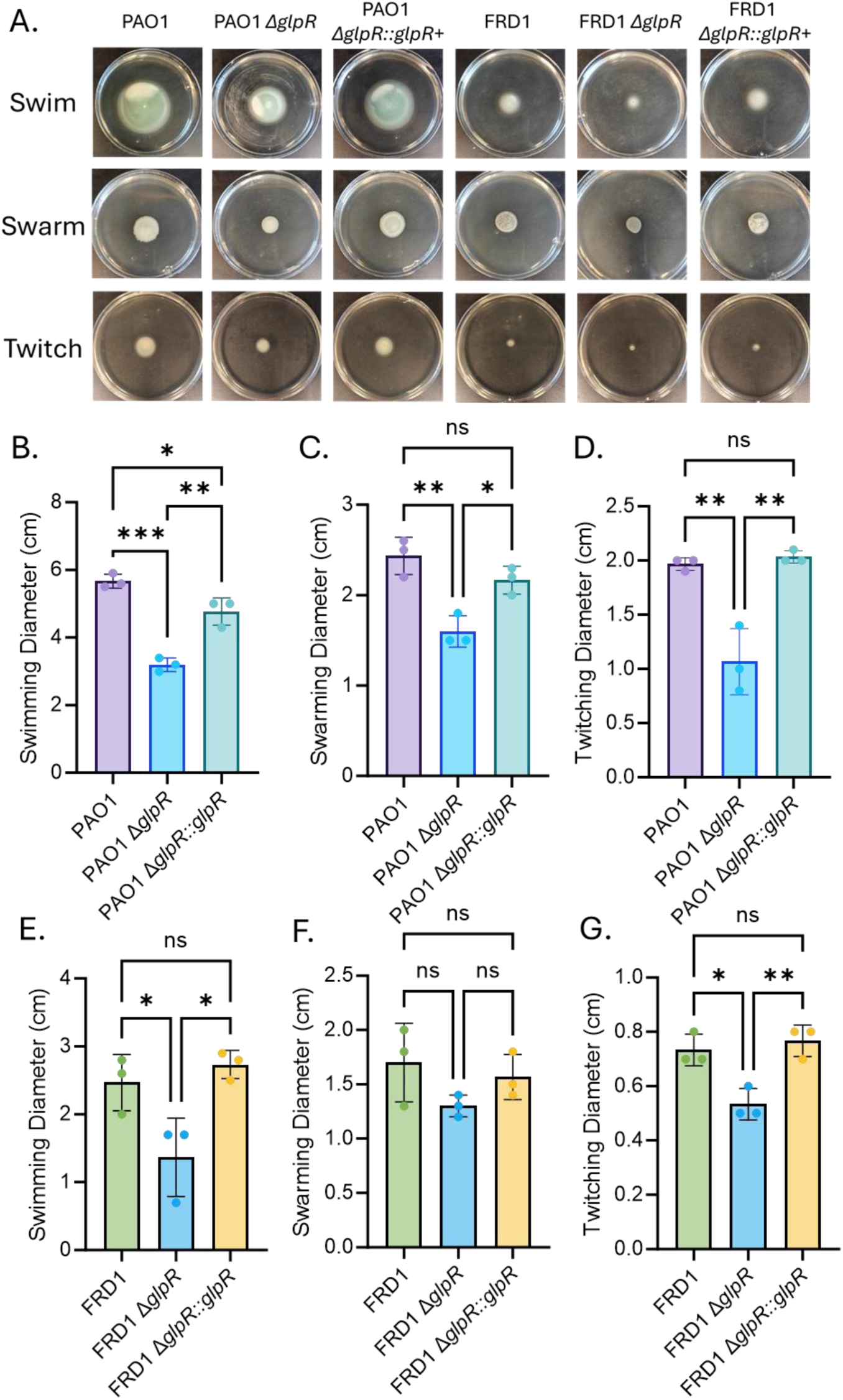
Loss of GlpR decreases motility of PAO1 and FRD1. A) Representative images of the swimming, swarming and twitching motility plates after incubating for 24 hours. B, C, D) Bar graphs showing the quantification of the swimming (B), swarming (C), and twitching (D) of the strains PAO1, PAO1 *ΔglpR*, and complemented PAO1. E, F, G) Quantification of the swimming (B), swarming (C), and twitching (D) of the strains FRD1, FRD1 *ΔglpR*, and complemented FRD1. (n=3) *Significance by one-way ANOVA. *P<0.05, **P<0.01, ***P<0.001

**Figure 4.**
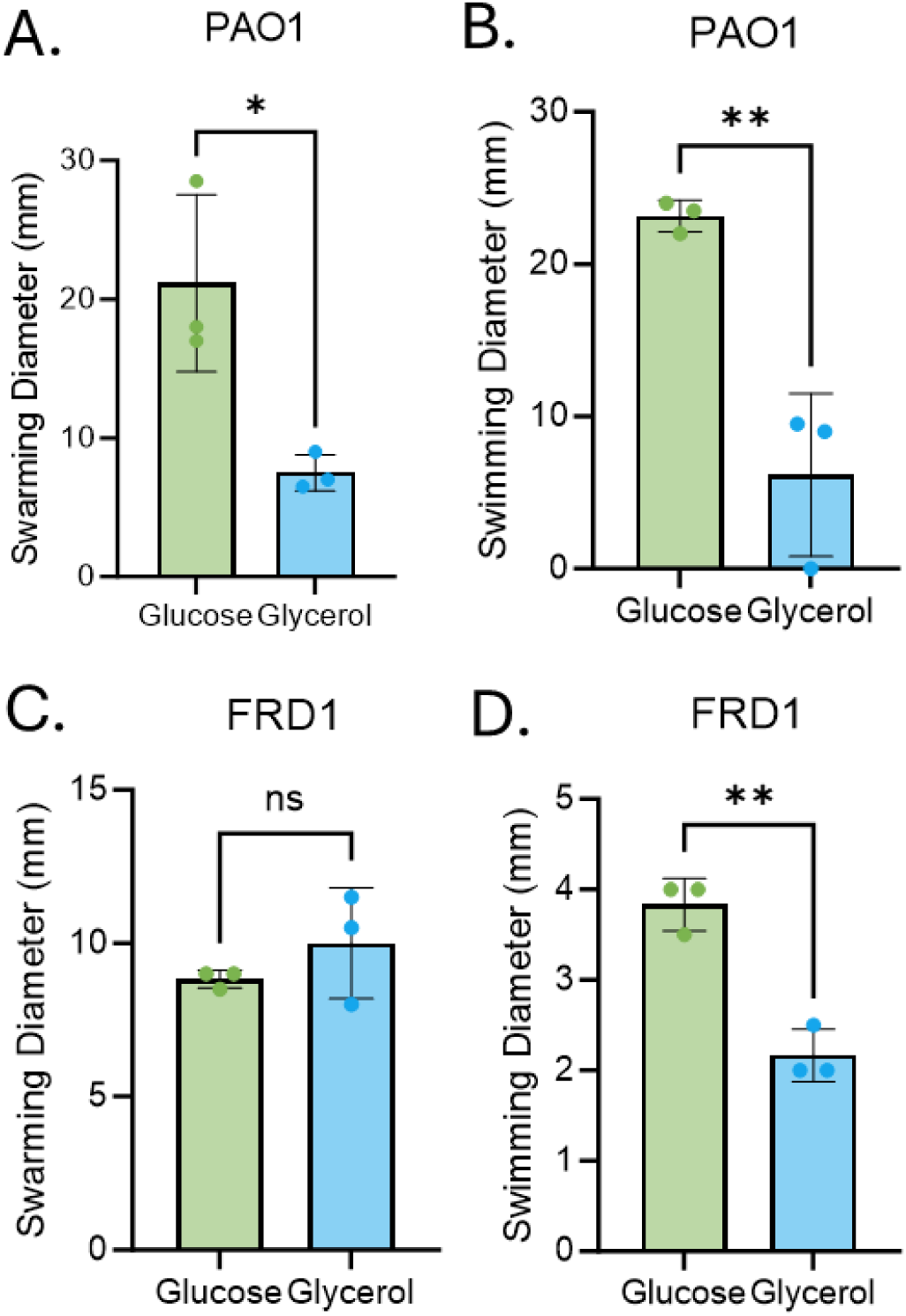
Growth on glycerol decreases swarming and swimming motility. A) Swarming motility results for PAO1 on minimal media with either 20 mM glucose or glycerol as the sole carbon source. B) Swimming motility of PAO1 grown on minimal media with glucose or glycerol. C) Swarming motility of FRD1 grown on minimal media with glucose or glycerol. D) Swimming motility of FRD’ on minimal media made with glucose or glycerol. (n=3) *Significance by t-test. *P<0.05, **P<0.01

In agreement with these findings, minimal media containing glycerol significantly reduced swarming (Figure 4A) and swimming (Figure 4B) motility of wildtype PAO1 compared to when grown on minimal media containing glucose. Additionally, while there was no change in the swarming of FRD1 when grown on glucose or glycerol (Figure 4C), swimming motility was significantly reduced in FRD1 on glycerol (Figure 4D). Taken together, our results support the conclusion that defects in GlpR reduce motility in both the PAO1 and FRD1 backgrounds and growth on glycerol impairs motility.

### Transcriptomics and in silico analysis reveal that GlpR potentially regulates genes outside of the canonical *glp* regulon

Our previous and current studies have determined that GlpR modulates *P. aeruginosa* biofilm development, motility, and virulence factor production. To dissect global changes caused by the loss of GlpR, we performed RNA sequencing analysis (27) and an *in silico* search to identify non-canonical genes that may be potentially regulated by GlpR. Loss of the GlpR repressor resulted in the upregulation of genes related to iron scavenging, fatty acid metabolism, glycerol metabolism, and phenazine production in the PAO1 background. In the iron scavenging group, the most upregulated gene was a hypothetical protein PA2427 (+3.4), followed by a transport component *fpvJ* (+2.9), a gene related pyoverdine synthesis *pvdG* (+2.9), and *foxR* (+2.6), which encodes an iron transport regulator. Differentially regulated genes related to fatty acid metabolism included a probable ABC transporter component PA2214 (+2.7), a fatty acid synthesis gene fabH2 (+2.1), a probable short chain dehydrogenase PA3330 (+1.9), and a probable FAD-dependent monooxygenase PA3328 (+1.7). As expected, we observed the upregulation of the *glp* regulon, including *glpD* (+4.5), the glycerol transporter *glpF* (+3.4), and a probable acyl-CoA thiolase PA3589 (+2.4) within the same region. For phenazine production related genes, one gene related to pyochelin transport was elevated (*fptB*, +4.4), in addition to three genes related to phenazine biosynthesis, *phzE1* (+2.8), *phzC1* (+2.3), and *phzG1* (+2.1). In addition, we observed the downregulation of many genes that encode for hypothetical proteins, including PA2186 (-3.7), PA2170 (-3.3) and PA2149 (-2.6). Further, we observed a downregulation of PA2933 (-3.9) a probable transporter, *morB* (-3.4) encoding a morphinone reductase, and *cifR* (-2.3), which encodes a transcriptional regulator (Figure 5A).

**Figure 5.**
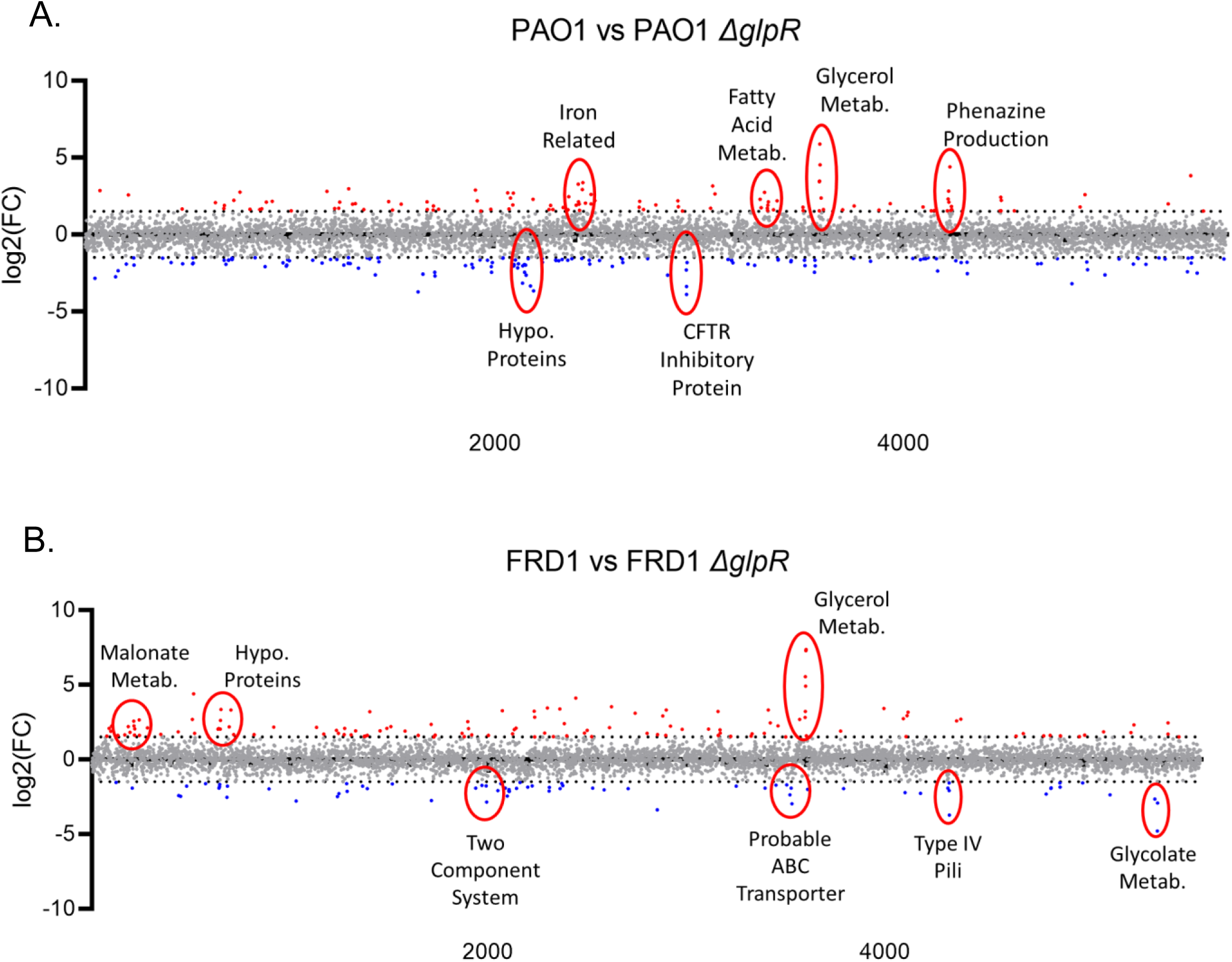
Clusters of differentially expressed genes from RNA sequencing analysis provide evidence for potential regulatory regions of GlpR. A) Scatterplot of all RNA sequencing results with highlighted clusters of up and down regulation for PAO1 Δ*glpR* compared to PAO1. B) RNA sequencing results for FRD1 Δ*glpR* compared to FRD1 with highlighted clusters of up and down regulation.

Differentially expressed clusters for FRD1 versus FRD1 *ΔglpR* included upregulated genes related to malonate metabolism, hypothetical proteins, and glycerol metabolism. The downregulated clusters included genes in a probable ABC transporter, type IV pili associated genes, and some glycolate metabolism genes. The most upregulated malonate metabolism genes were two genes encoding different subunits of malonate decarboxylase, *mdcD* (+2.6) and *mdcE* (+2.2) and also included the malonate transporter *madL* (+2.0). The hypothetical protein encoding genes included PA0644 (+2.0), PA0645 (+2.6), PA0648 (+3.3) and PA0698 (+3.3), which may encode a YbjN-like regulator. This group also contained *lapB* (+2.2), an alkaline phosphatase B that is annotated as being type 2 secretion system related. As with the PAO1 *ΔglpR* mutant, multiple glycerol metabolism genes were upregulated including *glpD* (+7.4), *glpM* (+7.3) which is necessary for alginate production, *glpF* (+5.5), and *glpK* (+4.9). Additionally, two genes upstream of the glycerol regulon were upregulated, including *ybaK* (+3.2) a tRNA deacylase and PA3579 which has been shown previously to be a glycerol kinase. The response regulator EraR was downregulated by -2.9-fold and is related to ethanol and methanol metabolism. The three most downregulated genes were the probable ABC transporter cluster PA3514 (-3.0), PA3504 (-2.4), and PA3510 (-1.9). Pilus associated genes were also downregulated, including the type IV pili group consisting of *flp* (-3.7) encoding a type IVb pilin, *rcpC* (-2.1) which is involved in pilus assembly, and PA4298 (-1.9), a hypothetical gene neighboring the type 2 secretion system operon. Lastly, two glycolate oxidase subunit genes were differentially expressed, *glcE* (-4.8) and *glcD* (-2.9) and this cluster also included a nearby gene PA5341 (-2.7) which may encode a hypothetical arginine transporter (Figure 5B). The differential expression of genes in the *glpR* mutant related to diverse functions compelled us to search for potential sites of GlpR. *In silico* analysis scanning the *P. aeruginosa* genome for potential GlpR binding sites, revealed the discovery of 456 putative binding sites. Not surprisingly, these genes included members of the *glp* regulon, but also genes involved in biofilm development, motility and chemotaxis, and drug resistance (Table 1), which support our previous and current findings (25, 27).

**Table 1.** GlpR predicted binding sites in PAO1 genome based on the consensus sequence. A selection of hypothetical binding sites for *P. aeruginosa* GlpR based on the published consensus sequence. Results were obtained using the motif search on pseudomonas.com and the sequence TWTWTYYSRM. These results consist of binding sites upstream of characterized genes and are grouped by functional area.

| Gene Name | Description |
| --- | --- |
| **Biofilm** |  |
| *bdlA* | Biofilm dispersion locus A <> hypo protein with DTW domain |
| *cdrA* | Cyclic diguanylate-regulated TPS partner A <> glycerate dehydrogenase |
| *cupA* | Fimbrial protein, adhesion, pilus, and oxygen response related operon |
| *cupE* | Pilin subunit (plus more cup regulon) |
| *lecA* | PA-IL like domain with galactose binding, related to cell-cell adhesion |
| *pslA* | Psl exopolysaccharide |
| *siaA* | Biofilm formation in response to carbon (CupA and RsmZ involved) |
| **Motility** |  |
| PA1442 | Flagellar genes (includes *fliMNOPQR-flhB* ) |
| PA1458 | Probable TCS for chemotaxis |
| PA1474 | Hypo flagellar hook length control protein <> heme exporter protein operon |
| *flgG* | Multiple flagellar genes after this site |
| *fliC* | Flagellar protein |
| **Glycerol acquisition** |  |
| *glpD* | Glycerol 3-phosphate dehydrogenase |
| *glpF* | Aminoacyl-tRNA editing, glycerol regulon |
| *glpT* | Glycerol 3-phosphate transporter |
| *plcH* | Hemolytic phospholipase C |
| **Drug Resistance** |  |
| PA0749 | Vancomycin resistance |
| PA4779 | hypo EamA like transporter (EmrE multidrug resistance efflux like) |
| *arnB* | Two Component System related to cationic antimicrobial peptide resistance and amino and nucleotide sugar metabolism |
| *mexP* | Multidrug efflux pump, lactoylglutathione lyase (pyruvate metabolism) |
| **Iron Scavenging** |  |
| *pvdR* | Efflux pump complex, pyoverdine related |
| *bfrB* | Bacterioferritin (siderophore) |
| *bphO* | Heme oxygenase |
| *pirA* | Siderophore transporter |

### Chromatin immunoprecipitation sequencing reveal*s* an expanded *GlpR* regulon

Based on our results supporting that the loss of GlpR affects the phenotypes of multiple characteristics affecting *P. aeruginosa’s* virulence, we aimed to determine whether the full regulon of GlpR may be expanded. We conducted chromatin immunoprecipitation sequencing (ChIP-seq) to identify GlpR binding sites, which identified 157 peaks shared by two biological replicates (Figure 6A). The top 100 genes associated with these peaks can be found in the supplemental excel file (S1) and include genes that influence motility, including *pilG, fliS, fliA, flp*, and *pilA*. Additionally, we observed peaks in alginate biosynthetic genes (*algD* and *amrZ*), which supports the increase in alginate we observed that influenced the viscoelasticity of FRD1 and FRD1 *ΔglpR.* Previously, we found that phenazine production, primarily pyocyanin was modified by loss of GlpR (27). ChIP-seq revealed binding sites in the phenazine genes *phzH, phzM,* and *phzB1*. Overall, the majority of the peaks mapped to hypothetical genes and genes involved in motility, drug resistance, regulation, phenazine and alginate production, as well as various cellular and housekeeping functions (Figure 6B).

**Figure 6.**
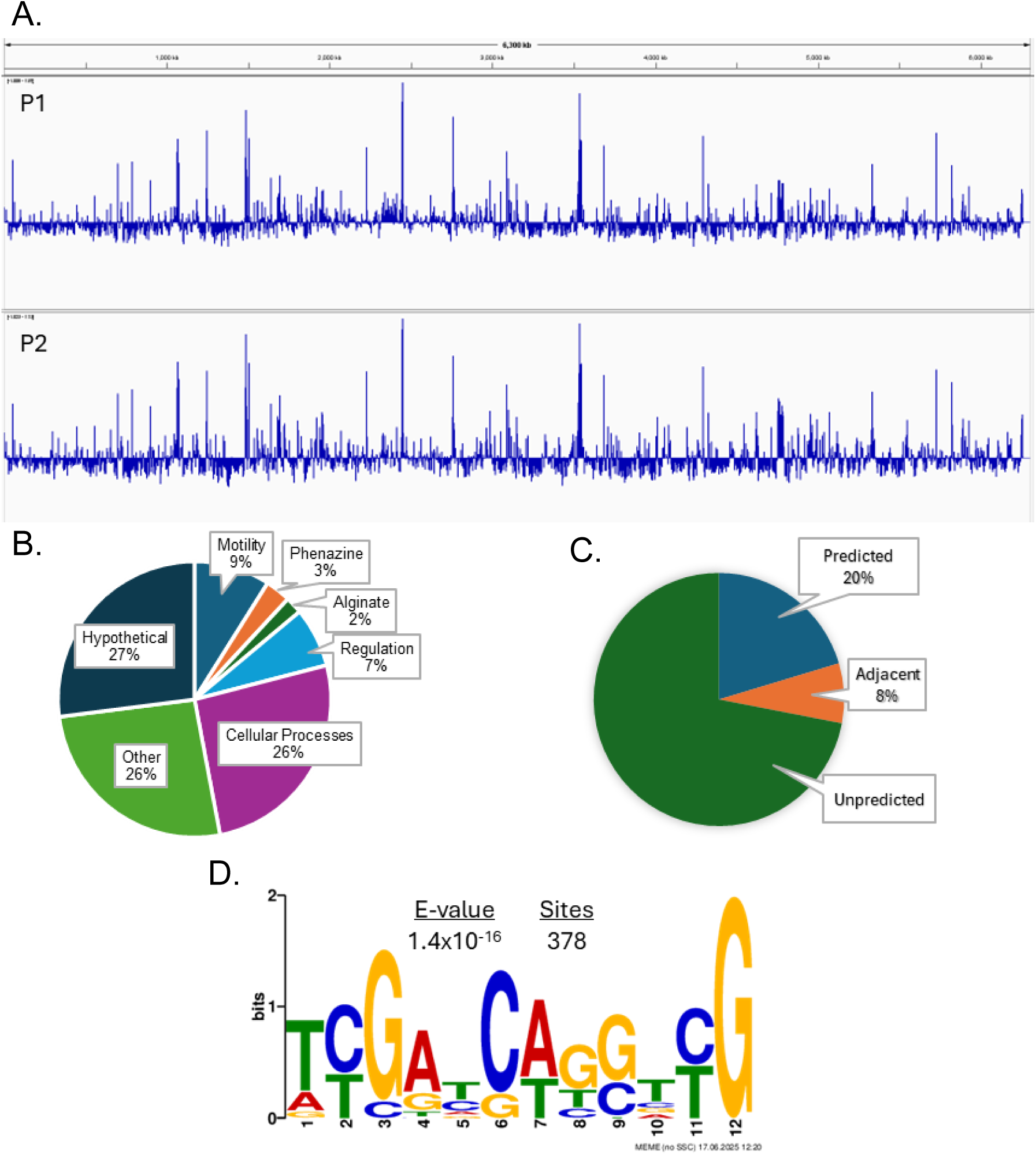
Chromatin immunoprecipitation sequencing reveals an expanded GlpR regulon. A) Visualization of the peaks found across the PAO1 genome for both biological replicates. B) A pie chart showing percentages based on the assigned category these peaks sort into. C) A pie chart of which peaks were predicted, adjacent to predicted genes or unpredicted based on our *in silico* search. D) The strongest binding motif generated based on the peaks we obtained.

Our prior *in silico* search was based on the previously published consensus sequence for GlpR, TWTWTYYSRM, which was derived from *E. soli’s* GlpR’s binding site. When comparing our peaks to the *in silico* search, only 20% were predicted, 8% were adjacent to a predicted gene and 72% were unpredicted (Figure 6C). Due to this expanded regulon, we used the MEME tool to compare the fasta sequences of the 157 peaks, to determine a potential motif for binding of GlpR in *P. aeruginosa*. The most prevalent binding site was TYGAYCWGSTYG which appeared 378 times in the shared peaks (Figure 6D).

## DISCUSSION

Nutrient availability and acquisition dictate the survival of microbes in the CF airway. As such, *P. aeruginosa* has evolved numerous strategies to acquire nutrients and adapt to the CF nutritional environment to facilitate its persistence within the complex CF airway(5, 16–18, 31, 32). Prior transcriptomics studies dissecting the metabolic activities of *P. aeruginosa* growing within CF sputum have revealed that glycerol metabolic genes are constitutively expressed in *P. aeruginosa* isolates recovered from this CF lung (23). These findings provided evidence that host-derived glycerol potentially mediates *P. aeruginosa* behavior and physiology. Our previous work illustrated that glycerol metabolism and defects in the *glp* regulon repressor, GlpR, induce biofilm development, antibiotic persistence, and the switch from the acute to chronic phenotype lifestyle of growth (25, 26). Building on these previous studies, we discovered that loss of the GlpR repressor decreased motility and increased the viscoelasticity of *P. aeruginosa* cultures grown in SCFM2, which are factors that promote biofilm development and reduce mucociliary clearance in the airway (33–35). Further, we provide evidence that GlpR regulates the expression of genes beyond the *glp* regulon loci. In sum, this study highlights new roles for GlpR that support its contribution to *P. aeruginosa* during adaptation of the CF airway.

The ability of a bacterium to sense and respond to metabolic fluxes is critically important for fitness. Although glucose is an abundant carbon source in the CF airway, glycerol is a preferred carbon source for clinical CF *P. aeruginosa* isolates (31, 36), suggesting that *P. aeruginosa* has undergone glycerol-dependent metabolic specialization. In our current study, loss of GlpR, and thereby constitutively expression of the glycerol metabolic genes decreased motility and increased the viscoelasticity properties of *P. aeruginosa* cultures. We hypothesize that the sensing, responding, and acquisition of host-derived glycerol serves as a nutritional cue for *P. aeruginosa* to switch from an acute to chronic lifestyle of growth, which includes reduced motility, biofilm formation, and antibiotic tolerance. We have previously reported that loss of GlpR increases the production of alginate (26), an exopolysaccharide responsible for the mucoid phenotype observed in many CF clinical isolates and is a hallmark of a chronic CF infection. We speculate that alginate-dependent increases in viscosity and elasticity in the FRD1 Δ*glpR* mutants support the persistence of *P. aeruginosa* in the airway. Increases in the viscosity or elasticity of *P. aeruginosa* cultures impede the penetration of antimicrobials or effectors of the host immune response, leading to recalcitrant biofilms, reduced susceptibility to antimicrobials, and reduced mucociliary clearance (33–35). Although highly effective modulator therapy (HEMT) with elexacaftor/tezacaftor/ivacaftor (ETI) has corrected some CFTR function and reduced the microbial burden (37–39), recent studies have shown that *P. aeruginosa* continues to persist after modulator therapy and has started to evolve in response to ETI treatment (11). We theorize that consistent availability of host-derived glycerol may drive *P. aeruginosa* persistence even with the advent of HEMT.

Transcriptomics and *in silico* analysis indicated that GlpR potentially regulates genes outside of the *glp* regulon, possibly through direct regulation or the reprogramming of carbon flux. To confirm the regulation of genes by GlpR outside of the *glp* regulon, we performed ChIP-seq to identify global GlpR binding sites. As expected, ChIP-seq identified several regions of GlpR binding in the *P. aeruginosa* genome. Remarkably, many of these regions included genes responsible for alginate biosynthesis, motility, and phenazine production, which directly support our previous findings that GlpR controls the expression of secreted virulence factors, alginate biosynthesis, motility, and biofilm development. It is important to note that one of the limitations of the ChIP-seq analysis is that it was only performed in the PAO1 (wound isolate) background and not the FRD1 (CF-adapted isolate) background, which are two genetically and phenotypically different strains. *P. aeruginosa* strains that live in the CF airway undergo microevolution over time to adapt to the conditions in the CF airway and much of this adaptation included the altered regulation of genes involved in virulence and carbon metabolism (11–15, 23, 31, 40–44). Differences between PAO1 and FRD1 in our transcriptomics analysis, as well as phenotypic assays, suggest that regulation of genes by GlpR are strain dependent, and this is likely due to adaptive pressures within the CF environment.

In summary, we report that GlpR functions as one of the modulators of *P. aeruginosa*’s physiology related to motility, viscoelasticity, iron scavenging, and biofilm formation. Through modifying the expression of genes necessary for alginate production, we observed increased alginate production previously that we have now linked with increased viscoelasticity of synthetic sputum (27). We also found that the loss of repression by GlpR, either through knockout or through growth on glycerol, leads to a decrease in motility for both PAO1 and FRD1. With transcriptomic evidence, GlpR works to influence key phenotypes of *P. aeruginosa* in response to its environment and the nutritional cues it detects. Further, the identification of the global GlpR regulon using ChIP-seq reveals that GlpR regulates genes that are outside of the canonical *glp* regulon. Taken together, our study reveals that GlpR may be an important transcriptional repressor that regulates the adaptation and microevolution of *P. aeruginosa* during a chronic CF infection.

## MATERIALS AND METHODS

### Bacterial strains and growth conditions

*P. aeruginosa* strains PAO1, PAO1 Δ*glpR*, FRD1, FRD1 Δ*glpR,* and complemented strains (25) were plated on Pseudomonas Isolation Agar and grown in liquid cultures in either lysogeny broth (LB) or cystic fibrosis synthetic sputum with mucin and DNA (SCFM2). All cultures were grown at 37°C. The transcriptomics data presented in this manuscript was adapted from a previous publication (26) and these data were deposited in Gene Expression Omnibus under accession number GSE280261.

### Swimming, Swarming, and Twitching Motility

All motility LB agar plates were made the morning of use and cooled to ambient temperatures for 3 hours prior to inoculation. For motility assessed on minimal media, 20mM glucose or glycerol was supplemented in 1x M9 salts. For swimming agar, 0.3% agar was used, and the plate was inoculated by dipping a 10 uL micropipette tip into overnight culture and piercing the agar plate in the center, without touching the bottom. For swarming plates, 0.6% agar was used, and plates were inoculated by adding 2.5 µL of overnight culture to the surface of the agar without touching it. All plates were then incubated for 24 hours in a humidified incubator at 37°C. After 24 hours, the diameter of the swimming and swarming motility was measured, and the plates were imaged. To measure twitching motility, we used 0.4% agar, however, the plates were inverted and allowed to dry overnight. Using a sterile 10 µL micropipette tip a colony was picked from the agar and then stabbed through the center of the plate to the plastic at the bottom, and the tip was jiggled up and down. The plates were then incubated overnight in a humidified 37°C incubator for 24 hours. The diameter of the halo of bacteria from twitching across the bottom of the plate was measured.

### Rheology Analysis

Rheology was conducted using the TA Instruments Waters Discovery Hybrid Rheometer. All samples were run with temperature soaking at 25°C with two technical replicates and three biological replicates per condition using 200 µL of culture used per replicate. Samples were generated by growing FRD1 or FRD1 Δ*glpR* in SCFM2 with or without 10 U/mL of alginate lyase overnight in 50 mL conical tubes. The tubes were grown at 37°C with shaking at 250 rpm.

### In silico search of predicted GlpR binding sites

An *in silico* search using the previously published GlpR consensus sequence, TWTWTYYSRM was conducted for potential binding sites on the Pseudomonas.com website. The DNA Motif search tool on pseudomonas.com was used with the PAO1 reference genome. Each location output was manually checked in the PAO1 genome for associated gene(s) and whether the site was intergenic or within a coding region. Sites located in potential promoter regions were noted.

### Construction of an Arabinose Inducible, FLAG-tagged GlpR

For Chromatin Immunoprecipitation Sequencing (ChIP-Seq) we generated a GlpR construct using the coding region and 50 bp of the upstream of the start site. A SacI restriction site was added to the 5’ upstream region and a HindIII site was added to the 3’ downstream region with a FLAG tag on the C terminal end. This fragment was then inserted into the multiple cloning site of the pHERD20T plasmid which contains an arabinose inducible promoter. The resulting plasmid was electroporated into TOP10 *Escherichia coli* cells and plated on LB containing 100 ug/mL ampicillin. Transformants were verified using PCR and restriction digest. The purified GlpR FLAG tagged pHERD20T construct was then transformed into the PAO1 *ΔglpR* and FRD1 *ΔglpR* using triparental mating. Transformed colonies were selected for using PIA plates with 100 ug/mL carbenicillin. Positive clones were verified with PCR analysis to confirm the presence of the GlpR-FLAG tagged construct.

### Chromatin Immunoprecipitation

PAO1 *ΔglpR*::GlpR-FLAG and FRD1 *ΔglpR*::GlpR-FLAG tags were grown overnight in LB +50 ug/mL carbenicillin at 37°C with shaking at 250 rpm. Overnight cultures were then subcultured by resuspending 1 mL of the overnight in 10 mL of LB +25 ug/mL carbenicillin and incubating them for 2 hours at 30°C with shaking. To induce, arabinose was then added at a final concentration of 0.2%, and the cultures were incubated at 28°C with shaking for 2 hours. After induction, the cultures were pelleted and resuspended in PBS + 1% paraformaldehyde and fixed for 10 minutes at room temperature. The fixative was then quenched with 2.5 M Glycine for a final concentration of 125 mM with agitation for 1 minute. The bacteria were pelleted at 5000 rpm for 5 minutes at 4°C then washed twice with ice cold PBS. For sonication and shearing, pellets were resuspended in PBS with protease inhibitor and were split into tubes of 300 uL each. Using a Bioruptor, samples were sonicated on low for 25 cycles, with 30 seconds on and 30 seconds off per cycle. The debris was pelleted with centrifugation at 7,000 rpm for 5 minutes and the supernatant was collected. The DNA concentration was checked by Nanodrop, and 100 ug of chromatin was used per IP reaction. The chromatin was diluted to 300 uL using ChIP Dilution Buffer consisting of 0.01% SDS, 1% Triton X-100, 1.2 mM EDTA, 16.7 mM Tris HCl, and 167 mM NaCl. 50 uL of the diluted chromatin was removed for input sample. To the remaining 250 uL, 1 ug of anti-FLAG antibody was added to the tube and incubated with rotating overnight at 4°C. The next morning, 10 uL of Protein A magnetic beads were added to each tube, and they were incubated at 4°C for 2.5 hours to bind. The beads were then washed 6x with 250 uL TBS and with 1X TE with each rotating at 4°C prior to being placed on the magnetic rack to remove the wash. The DNA was eluted by adding 100 uL of Elution Buffer (50 mM Tris-HCl, 10 mM EDTA and 1% SDS) and rotating 5 minutes at 4°C prior to collection. The resulting elution samples had NaCL added to a final concentration of 0.2M. The input samples were thawed, diluted to 200 uL with 150uL of elution buffer, then also had NaCl added to 0.2M. The input and elution samples were then incubated at 65°C for 4 hours to reverse crosslinking. The Zymo Research DNA Clean & Concentrate 5 kit was then used to clean up the DNA, and the samples were eluted in 30 uL of elution buffer from the kit. The DNA concentration of all samples was checked with a Nanodrop, and the eluted samples were further checked using Qubit due to their low concentrations. Samples were sent for sequencing and analysis to Genewiz.

For generating the peak graph We used the filtered bam files for our two ChIP-ed samples and their corresponding inputs to generate the graphs. The bigWig files were produced using the bamCompare tool in usegalaxy.com to simultaneously minimize background noise and generate a visualization of the peaks across the genome (45). The bigWig files were then viewed using the IGV software.

To determine the new motif based on the resulting peaks, we used the MEME tool on usegalaxy.com to search for the most common motifs (46). We set a limit of 10 motifs with a length of 5 to 20 base pairs. We enabled the search of the complementary strand and used the FASTA file generated from the bed file of the 157 merged peaks provided by Genewiz, pulling the sequences from the FASTA genome file for PAO1.

### Statistical analysis

All experiments were conducted using a minimum of 3 biological and 3 technical replicates. Unless otherwise noted, graphs represent sample means ± standard error of the mean (SEM). Analysis of variance (ANOVA) or t-test statistical tests were performed using GraphPad Prism 10 (San Diego, CA).

## Acknowledgements

This work was supported by grants and funds awarded to J.A.S from the National Institute of General Medical Sciences (R35GM142748), the UAB Heersink School of Medicine Pittman Scholars Fund, and start-up funds from the UAB Department of Microbiology.

